# Looking for a reference for large datasets: relative reliability of visual and automatic sleep scoring

**DOI:** 10.1101/576090

**Authors:** C. Berthomier, V. Muto, C. Schmidt, G. Vandewalle, M. Jaspar, J. Devillers, G. Gaggioni, S. L. Chellappa, C. Meyer, C. Phillips, E. Salmon, P. Berthomier, J. Prado, O. Benoit, M. Brandewinder, J. Mattout, J. Maquet

## Abstract

**Study Objectives:** New challenges in sleep science require to describe fine grain phenomena or to deal with large datasets. Beside the human resource challenge of scoring huge datasets, the inter- and intra-expert variability may also reduce the sensitivity of such studies. Searching for a way to disentangle the variability induced by the scoring method from the actual variability in the data, visual and automatic sleep scorings of healthy individuals were examined.

**Methods:** A first dataset (DS1, 4 recordings) scored by 6 experts plus an autoscoring algorithm was used to characterize inter-scoring variability. A second dataset (DS2, 88 recordings) scored a few weeks later was used to investigate intra-expert variability. Percentage agreements and Conger’s kappa were derived from epoch-by-epoch comparisons on pairwise, consensus and majority scorings.

**Results:** On DS1 the number of epochs of agreement decreased when the number of expert increased, in both majority and consensus scoring, where agreement ranged from 86% (pairwise) to 69% (all experts). Adding autoscoring to visual scorings changed the kappa value from 0.81 to 0.79. Agreement between expert consensus and autoscoring was 93%. On DS2 intra-expert variability was evidenced by the kappa systematic decrease between autoscoring and each single expert between datasets (0.75 to 0.70).

**Conclusions:** Visual scoring induces inter- and intra-expert variability, which is difficult to address especially in big data studies. When proven to be reliable and if perfectly reproducible, autoscoring methods can cope with intra-scorer variability making them a sensible option when dealing with large datasets.

**Statement of Significance:** We confirmed and extended previous findings highlighting the intra- and inter-expert variability in visual sleep scoring. On large datasets those variability issues cannot be completely addressed by neither practical nor statistical solutions such as group training, majority or consensus scoring.

When an automated scoring method can be proven to be as reasonably imperfect as visual scoring but perfectly reproducible, it can serve as a reliable scoring reference for sleep studies.

## Introduction

Sleep studies are increasingly demanding and require fine grain characterization of sleep in bigger data sets,^1, 2^ collected in large population cohorts for phenotypic,^3, 4^ longitudinal,^5^ multi-center,^6^ or epidemiologic studies.^7^

Visual sleep scoring is the gold standard for sleep analysis. The typical output of sleep analysis is the succession of sleep stages across time, the hypnogram, resulting from the visual identification and classification by an expert applying somewhat arbitrary although consensual criteria.^8, 9^

However, visual scoring is affected by inter and intra-expert variability as well as inter-site variability.^10–14^ This results from the difficulty for different human experts to achieve exactly the same scoring or for a single expert to obtain the exact same scoring over time, for a given recording.^3, 15–18^

Training sessions partially alleviate this issue and improve the homogeneous application of the scoring rules.^15, 19^ However, the benefits of this strategy are limited to the period of training. To prevent a scoring “drift” over time, training sessions must be repeated.^1^ In the daily practice, this countermeasure does not cope with possible time-on-task effects often observed in monotonous tasks,^20^ especially when confronted with large datasets. Furthermore, periodic reappraisal of scoring rules^16–18, 21^ adds to the difficulty of applying consistently these rules, adding another source of variability.^19^ In conclusion, visual scoring introduces exogenous, potentially confounding, sources of variability. Mitigating solutions can be implemented only when dealing with a “reasonable” amount of data.

In order to score datasets made of thousands of sleep recordings, an error-minimizing, stable and reproducible method must be considered. Although such automatic method would not sizeably reduce inter-scorer variability, which is largely due to epochs on which scoring rules application is ambiguous^12, 22^, it would resolve intra-scorer variability.

In this study visual and automated scorings of healthy individual recordings were examined with respect to inter- and intra-expert variability, using a validated automatic scoring algorithm, Aseega.^23^

## Methods

### Participants and Experimental design

Data were extracted from a study aiming at characterizing sleep-wake related phenotypes. The study was approved by the Ethics Committee of the Faculty of Medicine of the University of Liège and participants gave their written informed consent. All volunteers were free of medication or psychoactive drugs, non-smokers and moderate caffeine and alcohol consumers. Semi-structured interviews established the absence of medical, traumatic, psychiatric or sleep disorders. Exclusion criteria were a poor sleep quality as assessed by the *Pittsburgh Sleep Quality Index* (score >5),^24^ excessive daytime sleepiness (score >10 on Epworth Sleepiness Scale),^25^ night shifts during the preceding year, travels through more than one time zone during the last three months, depression (score >19 on the Beck Depression Inventory II)^26^ or a body mass index >27 kg/m^2^.

The experimental design consisted of a 3-week actigraphic field recording followed by a 6-day laboratory protocol. A first habituation night aimed at ruling out participants with sleep disorders. Then four full-night polysomnographic (PSG) recordings were acquired under successive sleep conditions: 8h baseline night (BAS), extended sleep opportunity (EXT, i.e., 12h sleep opportunity at night, and 4h nap in the afternoon), 8h night preceding 40h of sleep deprivation (BEF) and a recovery night (REC). Sleep schedule was individually adjusted to habitual sleep time, as assessed during the pre-laboratory field study.

Sleep EEG data were recorded using a V-Amp 16 amplifier (Brain Products GmbH, Gilching Germany). EEG data were digitized at a sampling rate of 500 Hz with a bandpass filter from DC to Nyquist frequency and magnitude resolution of 0.049 μV/bit. During BAS, EXT, BEF and REC nights, recordings included 10 EEG channels (F3, Fz, F4, C3, Cz, C4, Pz, O1, O2, A1, reference: right mastoid), two EOG (horizontal and vertical EOG), two chin EMG and two ECG.

For the current retrospective study, the recording database includes data collected in 24 healthy, young, male participants (aged 21.6 ± 2.5 years, in the range 18-26), randomly selected from a larger sample of male volunteers.

### Automatic Scoring

Aseega is a comprehensive method for automatic sleep scoring, based on the analysis of the single EEG bipolar signal Cz-Pz.^23^ The automated procedure comprises 3 steps: preprocessing, analysis, and classification. The basic principle of the algorithm in each step is to use data-driven instead of *a priori* thresholds. This allows the algorithm to cope with major sources of variabilities such as individual characteristics of sleep EEG, recording conditions and sleep conditions, without using any prior information regarding the expected night contains. The algorithm is designed so as to be totally deterministic and reproducible, i.e. the results derive from the application of constant predefined rules leaving no place for stochastic decision nor evolution over time: a given version of the algorithm running x time on the same recording will produce exactly the same result each time. The main outputs of the algorithm are the hypnogram with spectral analysis and sleep microstructures including spindles and alpha bursts. Aseega typically completes the analysis of an 8h-night in about 5 minutes, on a standard PC. This value may slightly increase or decrease (± 1min) with the number of detected microstructures.

Performances of an earlier version (v1.3.14) were validated in a first cohort of healthy subjects.^23^ A newer version (v3.19.36) of the algorithm was used in the present study. The current version was designed to be able to cope with patient or ambulatory recordings thanks to improved identification of wakefulness and enhanced robustness to signal artefacts (movement artefacts, loose sensors, electromagnetic pollution), resulting simultaneously in more accurate feature extraction, dynamic threshold settings and sleep stage classification by the algorithm.

Interestingly, these improvements also refined the analysis performed in healthy participants, tested in laboratory conditions. This is shown by comparing the performance of both versions of Aseega, the one published in 2007^23^ and the current one, on the 2007 dataset, following the same procedure and using the same criteria as in 2007 (see table 1).

**Table 1:**
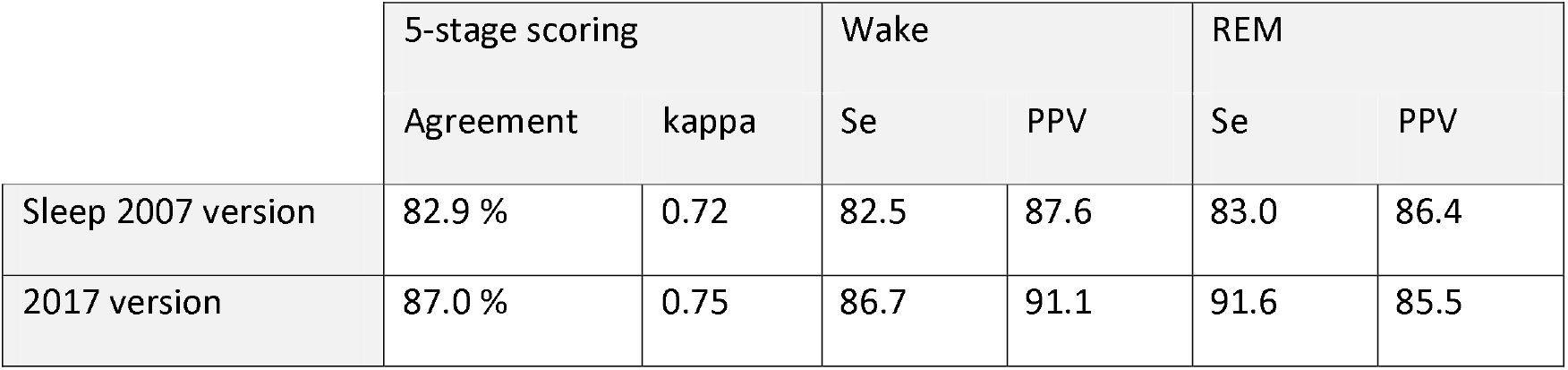
Comparison of the previous (2007) and current automatic scoring version when analyzing the dataset of the 1^st^ 2007 validation.

### Visual Scoring

Six expert visual scorers, five seniors and a junior, from the same sleep laboratory were trained together to homogenize the application of the AASM scoring visual rules^9^ on 30s-epochs, using the same scoring software, FASST, (http://www.montefiore.ulg.ac.be/~phillips/FASST.html).^27^ The training was considered achieved for a given expert when 80% of scoring agreement was reached with the expert leader, considered as the laboratory gold standard. Concerning the specific case when technical artefacts obscured the EEG, since the study framework was sleep research, such epochs were labeled as artefact.

In this paper, a human scorer is designated by ‘visual scorer’, ‘V’, or ‘expert’ while ‘A’ refers to automated analysis Aseega. Depending on the context, ‘scorer’ and ‘scoring’ refer to automated or visual scorer.

### Scoring variabilities

The inter-scorer variability is defined as the difference that can be measured between the scorings of two or more scorers, and raises no ambiguity as for its assessment. The consensus scoring refers to the labels on which all scorers involved, two or more according to the situation, agreed. In this study, the consensus is determined by statistical means, not as the result of verbal discussion between experts.

The intra-scorer variability is defined as the evolution in the way to score of a given scorer when compared with a reference. It can be tackled in two distinct ways according to the reference that is chosen. A first approach is to use the scorer as its own reference, the typical score-rescore agreement.^28^ A second approach is to use a confirmed sleep expert, supposed to be stable over time, as the scoring reference; the variability over time of the inter-scorer agreement (tested scorer vs. sleep expert) is then interpreted as the intra-scorer variability of the tested scorer. This corresponds to the typical training situation, where the increase over time of the inter-scorer agreement (trainee vs. sleep expert) is interpreted as the intra-trainee variability and is taken as a progress of the trainee.^15, 29^

As the used algorithm is reproducible, we propose a third approach to assess intra-expert variability, similar to the second one, where a validated autoscoring method is used as the reference to assess fluctuations in visual scoring.

### Datasets

The recording database consisted in two datasets, DS1 and DS2, referred to as scoring condition, which differed according to the temporal proximity to the training sessions. The scoring sessions of DS1 started 1.5 month after DS1 within a 6 months period. DS1 and DS2 excluded the recordings used for the training sessions.

The first dataset includes 4 recordings from 2 volunteers: 3 recordings from subject #0981 (BAS, EXT, and BEF) and 1 from subject #1004 (BAS). Each recording was scored independently by all 6 experts, thus providing 24 visual scorings, and by ASEEGA, providing 4 automatic scorings.

DS1 was used to assess the **inter-scorer** agreements as follows:

- by computing the inter-expert (V) agreement;
- by computing the inter-scorer (V+A) agreement, pooling visual and automatic scorings;
- by comparing automatic (A) scoring vs. visual (V) scorings.

The second dataset includes the 4 sleep nights (BAS, EXT, BEF and REC) from 22 other subjects, providing 88 recordings. Each recording of DS2 was scored twice: once visually by one of the 6 experts, providing 88 visual scorings, and once by ASEEGA, providing 88 automatic scorings. Note that when a subject was assigned to an expert, the 4 nights of this subject were scored by this expert.

Based on the automatic-expert agreements, DS2 was used to assess the **intra-expert** agreement as follows:

- by comparing DS1 vs. DS2 to assess the impact of the scoring condition;
- by comparing BAS, EXT, BEF, REC to assess the impact of the sleep condition.

The composition of the two datasets is summarized in Table 2.

**Table 2:**
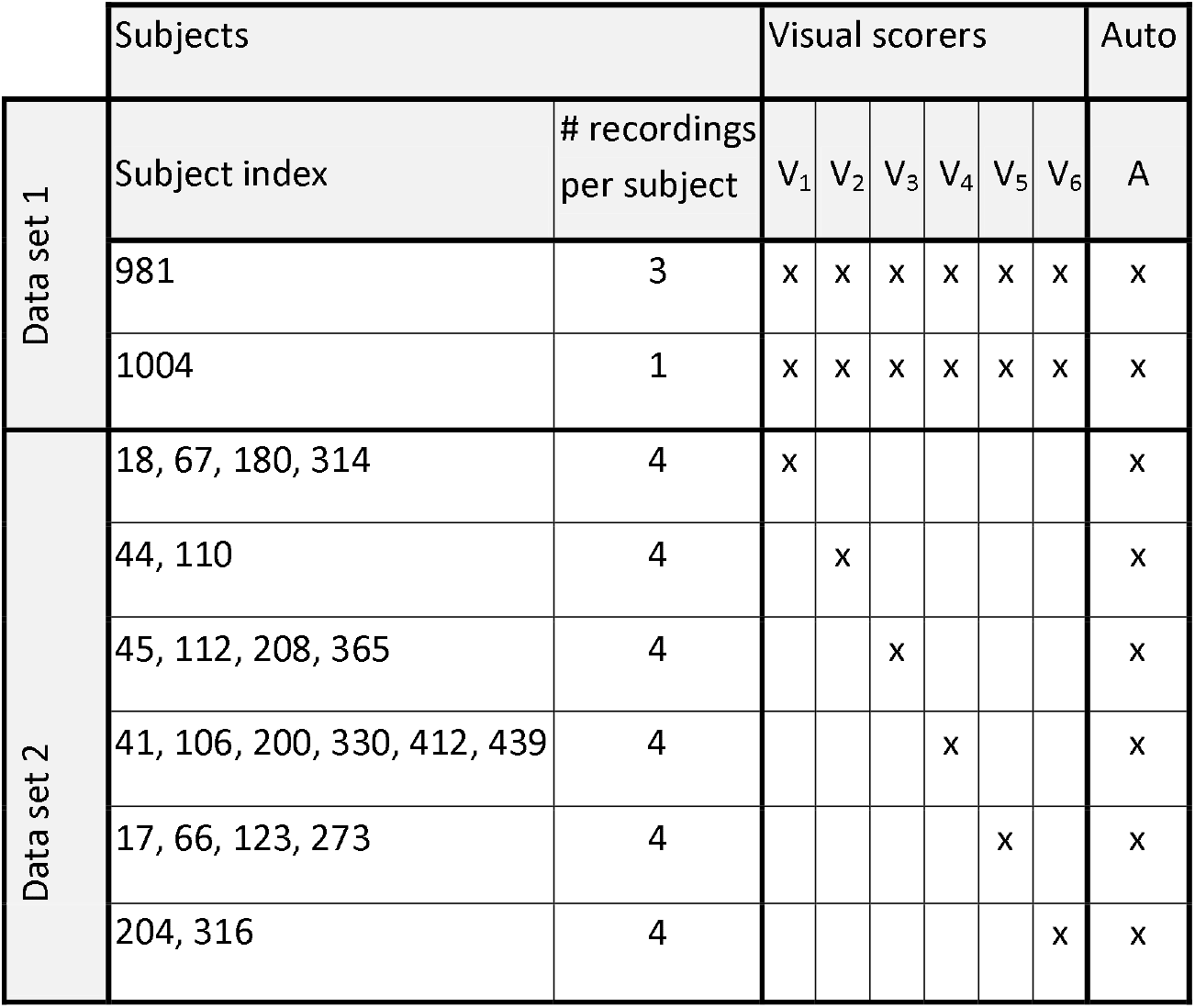
Cohort breakdown into 2 data sets DS1 & DS2, according to the number of times recordings were visually scored by the 6 visual scorers involved (V_1_ …V_6_). Each subject may provide for 1 to 4 nights. All scorers scored all recordings of DS1 (4 nights). In DS2, subjects were assigned to scorers who scored the 4 nights of such subject. In other words, each recording of DS1 was visually scored 6 times, whereas each recording of DS2 was visually scored only once.

### Statistics

Scorings were compared on an epoch-by-epoch basis. In order to mitigate the risk of overweighting the agreement calculated on short night and underweighting the agreement calculated on long nights, comparisons in each dataset were based on pooled night scorings rather than on averaged agreements over nights.^23^ Hence, for each dataset separately, the hypnograms of a given scorer were concatenated into a single continuous sequence of sleep stages.

After the computation of the contingency matrix, epochs labeled “artefact” by Aseega or by visual scoring were discarded from subsequent statistical analysis.

Two metrics of epoch-by-epoch agreement between scorers, whether automatic or visual, were used. First, the percentage agreement, defined as the percentage of epochs that were assigned the same label, i.e. sleep stage, by 2 or more scorers. Second, the Conger’s kappa (κ), which is the generalization of the Cohen’s kappa coefficient^30^ to the comparison of more than two raters. In the case of 2 raters, Conger’s kappa and Cohen’s kappa are equivalent.^31^ In order to have an homogeneous statistical criteria we computed Conger’s kappa instead of Fleiss kappa, sometimes used in multi-scorer comparison^19^ but which reduces to a different 2-rater agreement coefficient called Pi.^32, 33^

### Dataset 1

In a first step, we studied visual scoring in the light of the number of experts involved in the process. Do experts converge and how? How adding up experts does impact on inter-expert agreement? We studied the evolutions of both the visual percentage agreement and kappa as functions of the number of experts involved, N_e_. For a given N_e_ (from 2 to 6), all possible combinations N_c_ of experts were computed, providing distributions of agreements.

In a second step, we added autoscoring to the pool of visual scorings and assessed the inter-scorer agreement. We performed pairwise comparisons using both kappa and percentage agreement, with every possible pairs of scorings (V and A). We also performed comparisons between autoscoring and visual full consensus V_all_, as well as between each visual scoring V_i_ and the consensus of all the other experts (V_exc.j_, **i≠j**).

In a third step, we compared all scorers (V and A) with respect to their individual contribution to the overall agreement, questioning whether automated analysis contributes differently to the overall agreement than visual experts, and if so how. A global kappa, κ_G1_, was computed on all 7 scorings available (6 visuals + A). Partial kappa coefficients were then computed for all possible pools of 6 scorings where only 1 scorer was left out of the pool (pool minus scorer 1, pool minus scorer 2, etc.).

The specific case where autoscoring was removed stands for an assessment of the impact of adding automated analysis to the pool of experts.

The 6 experts are expected to show a very high agreement after the training sessions, thus minimizing the weight of the seventh (automated) scoring on the global kappa κ_G1_. To enhance what could be the impact of a bad autoscoring, we introduced a synthetic artificially altered automated scoring. We simulated two different random scorings: one where sleep stages have the same prevalence, but labels were randomly shuffled and one where labels themselves were randomly generated regardless of the true sleep stage prevalence. We refer to those scorings as “randomly shuffled autoscoring” and “fully random autoscoring”, respectively. In order to test for significance, 1000 different randomly shuffled and 1000 different fully random autoscorings were generated. For each random generation the global kappa was calculated for the 6 experts plus each altered automated scoring, providing a distribution of global kappa coefficients.

As an alternative to consensus scoring, which is a harsh approach since any epoch where only one expert disagrees is rejected, the fourth step consisted in replacing the visual full consensus scoring by the visual majority scoring, V_Maj_.^15^ For each epoch, we considered as V_Maj_ the scoring decision which brings together at least N_Maj_ experts, with N_Maj_ varying from 2 to 6. Note that the case of a majority of at least N_Maj_=6 experts is equivalent to the full consensus case. In case of ex aequo majority scorings, e.g. REM decision for three experts and N2 decision for the other three experts for a given epoch, the epochs were considered as non-valid and discarded when computing the percentage agreement. Likewise, when an epoch did not meet the minimum number of majority members requested, e.g. REM decision for three experts and N2 decision for three other experts whereas the minimum number of experts requested to constitute a majority is N_Maj_=4. In order to investigate the link between the experts and the majority scoring, the comparisons V_i_ vs. V_Maj_ were also performed for each expert V_i_ and each N_Maj_.

In a fifth and last step, since the visual full consensus is the most reliable visual reference as all epochs raising doubts are rejected, the contingency matrix between automatic scoring and the visual full consensus were computed to provide Sensitivity (Se) and Positive Predictive Value (PPV) for all sleep stages.

### Dataset 2

In this dataset, each recording was scored by only one of the six experts (see Table 2).

In a first preliminary step, the global kappa, κ_G2_, and the percentage agreement between automatic and visual scorings were computed. Note that since DS2 can only provide pairwise comparisons κ_G2_ cannot be compared with κ_G1_. The recordings scored by a given expert were then pooled together to provide for each expert a pairwise comparison with autoscoring.

The second step aimed at assessing the intra-expert variability according to the scoring condition, i.e. across datasets. Is visual scoring impacted by the scoring condition and how? We used autoscoring as reference since it is blind to the scoring condition. The auto-visual agreements on DS2 were compared with the corresponding pairwise agreements obtained on DS1.

A third step was to demonstrate that the comparison of auto-visual agreement between DS1 and DS2 was not biased by the difference in size of the datasets. We computed the agreement on random subsamples of DS2 data, each subsample having the same number of epochs as in DS1. 1000 subsamples of DS2 were randomly computed to provide a distribution of kappa coefficients and percentage agreements.

The fourth step aimed at assessing the intra-expert variability according to the sleep condition (BAS, EXT, BEF and REC). We used again autoscoring as reference, since it is also blind to the sleep condition. Kappa coefficients and percentage agreement were computed for each expert and in each sleep condition.

Finally, as a fifth and last step, the contingency matrix yielded Se and PPV for all sleep stages, without distinguishing between experts.

## Results

### Dataset 1

The figure 1 shows the qualitative comparison between all 7 scorings on night EXT night of subject #0981.

**Figure 1.**
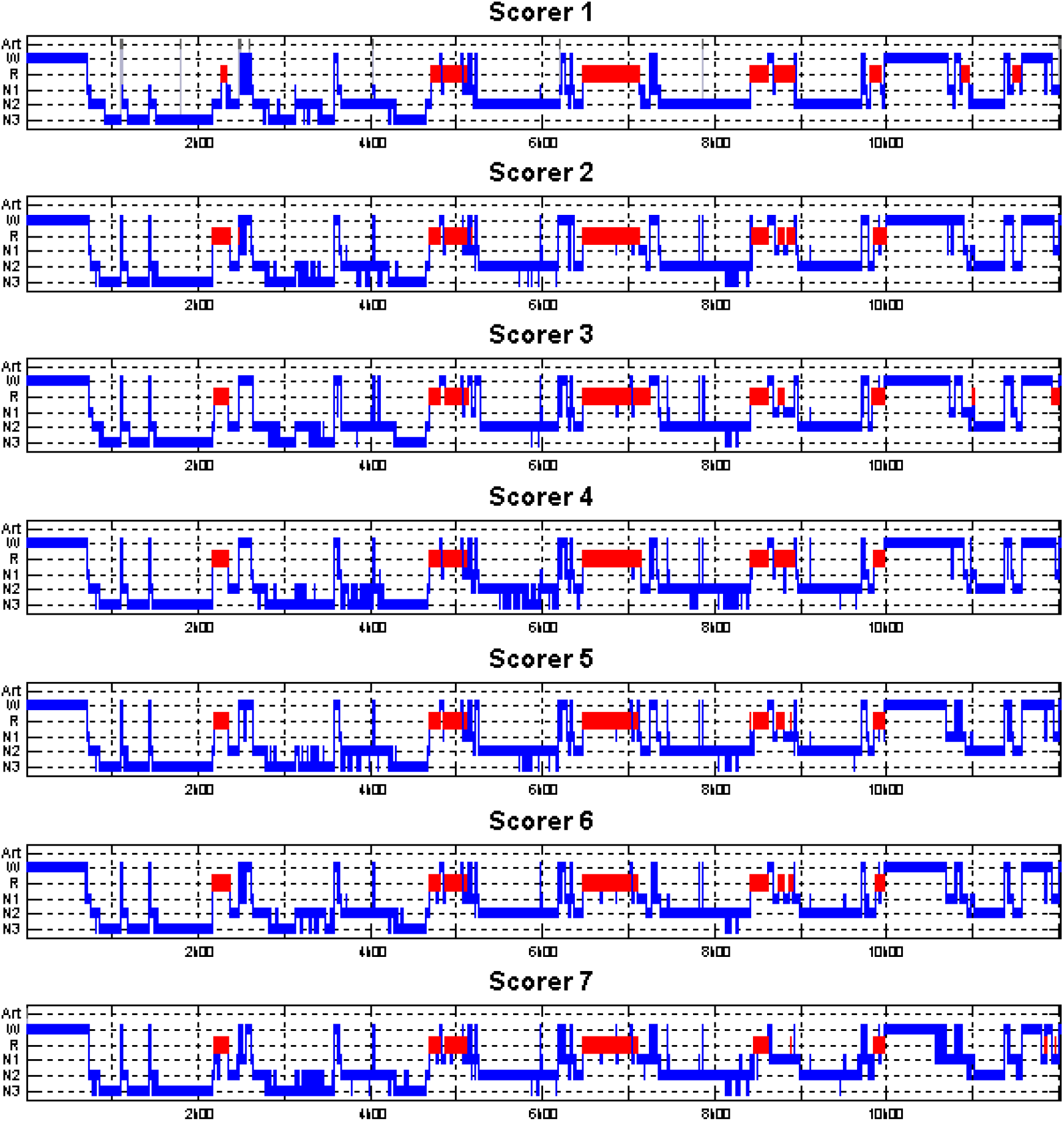
Representative 5-state hypnograms obtained using automatic scoring (scorer 1) and 6 independent experts (scorers 2 to 7) on one recording of DS1. On the y-axis are reported the various sleep stages: Wakefulness (W), Rapid Eye Movement Sleep (R), NREM sleep stages 1 to 3 (N1, N2, N3). The artefact (Art) label stands for the specific case when technical artefacts obscured the EEG. The recording time is reported on the x-axis.

Out of 4,384 epochs, 39 (0.89%) were classified as artefact by the automatic scoring versus 38 epochs (0.87%) by the experts. However, the number of epochs labeled as artefact varied substantially between experts: 0, 0, 1, 1, 33 and 3 respectively. Moreover, none of these epochs were identical across scorers, while only one epoch was in common between the automatic and the visual classifications. We thus discarded a total number of 76 (1.7%) epochs, leaving 4,308 clean epochs for subsequent statistical analysis.

In the first step, the inter-expert agreement as a function of the number of experts N_e_ (fig. 2) showed that the larger the number of experts, the lower percentage agreement and kappa variance, while mean kappa remained stable. The percentage agreement across all six visual scorers was 68.7% and the corresponding kappa was 0.81.

**Figure 2.**
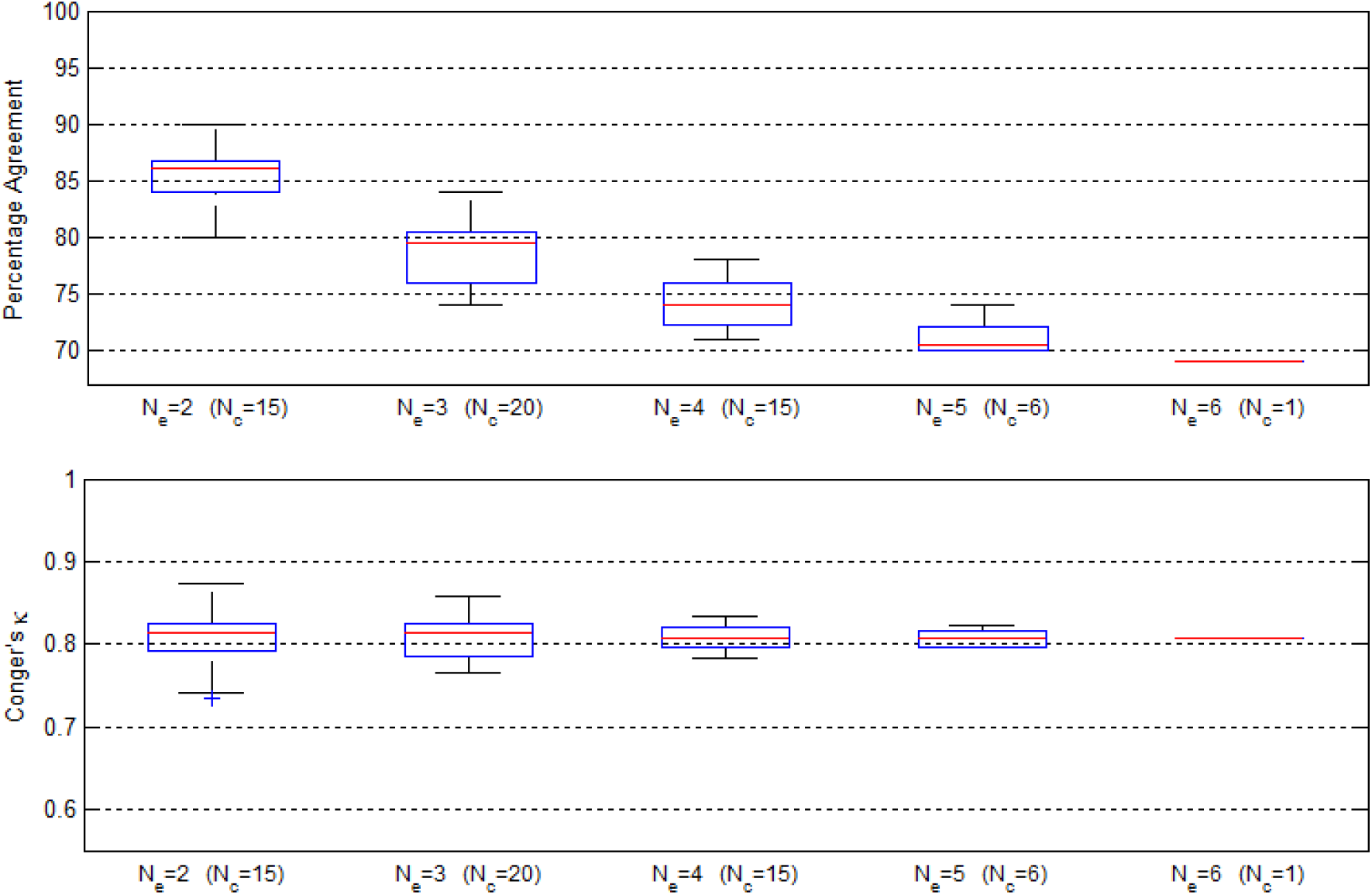
Inter-expert agreement: evolution of the agreement according to the number of visual scorers involved. The consensus agreement (percentage agreement, upper plot) and the inter-expert agreement (Conger’s kappa, lower plot) are drawn according to the number of experts N_e_ included in the pool. All the expert combinations N_c_ have been computed (N_c_=15 for agreement between N_e_=2 experts out of 6, N_c_=20 for 3 experts out of 6, etc.) for each possible number of experts (N_e_ = 2 to 6), yielding distributions of agreements. Note that for N_e_=6 experts, the number of possible combinations is reduced to N_c_ = 1.

Concerning the inter-scorer variability ([min − max], μ=mean), the 6 pairwise kappa coefficients between autoscoring and each scorer separately, A vs. V_i_, ranged [0.72 − 0.79], μ=0.75, and the 15 pairwise kappa coefficients between visual scorers, V_i_ vs. V_j_, ranged [0.73 − 0.87], μ=0.81 (fig. 3a). The corresponding percentage agreements A vs. V_i_ ranged [79.1% − 84.7%], μ=81.7%, when V_i_ vs. V_j_ ranged [79.8%−90.5%], μ=85.6% (fig. 3b).

**Figure 3.**
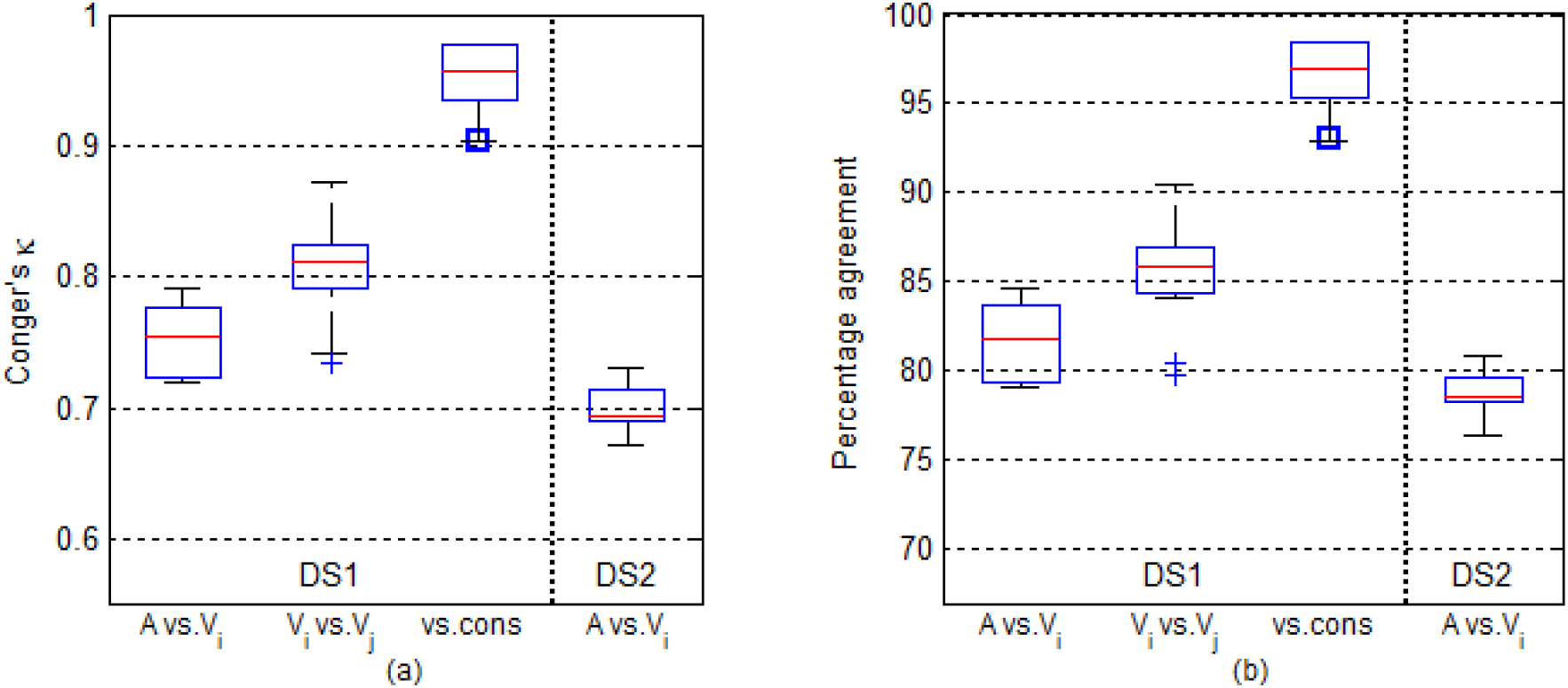
Inter-scorer and intra-scorer variability, pairwise comparisons. Comparisons are presented via (a) Conger’s κ and (b) percentage agreement. These 2 graphs are divided into results on DS1 (the 3 boxplots on the left) and on DS2 (4^th^ box on the right). The comparison between automatic scoring and each visual scorer (A vs. V_i_) is represented on the 1^st^ plot, the comparison between each pair of visual scorers (V_i_ vs. V_j_, with i≠j) is represented on the 2^nd^ plot. On the 3^rd^ plot, are reported the agreements between each visual scoring and the scoring consensus built by the other experts. The specific case of the agreement between autoscoring and the full visual consensus is highlighted in blue square. The 4^th^ boxplot on the right represents the A vs. V_i_ pairwise comparison on DS2. For each of the 2 graphs, the 3 plots on the left illustrate the inter-scorer variability on DS1 data and confirm that comparisons with scoring consensus provide far better agreements since the doubtful epochs are discarded. The left-most plot, together with the right-most plot, illustrate the intra-scorer variability between datasets (scoring condition, i.e. temporal proximity with the training sessions), using automatic analysis as the reference.

As expected, pairwise comparisons yielded lower agreements than comparisons involving consensus scoring since the latter discards ambiguous epochs. Accordingly, the kappa between A and the visual full consensus V_all_ was 0.91, whereas kappa between each individual expert and the consensus of the other experts ranged [0.90 − 0.98], μ=0.95 (fig. 3a, 3^rd^ boxplot). Similarly, the percentage agreement was 93.1% between A and V_all_, and [92.9% − 98.5%], μ=96.5% between each V_i_ and the consensus of the other visual scorings (fig. 3b, 3^rd^ boxplot).

The global kappa over DS1 obtained by comparing all scorings (V+A) was κ_G1_ = 0.79. It is barely lower than the kappa obtained by comparing visual scorings (V) only (fig. 4a). By contrast the two simulated random scorings proved to be highly deleterious, mean κ = 0.58 (fig. 4b) and 0.56 (fig. 4c) for the randomly shuffled and fully random scoring respectively.

**Figure 4.**
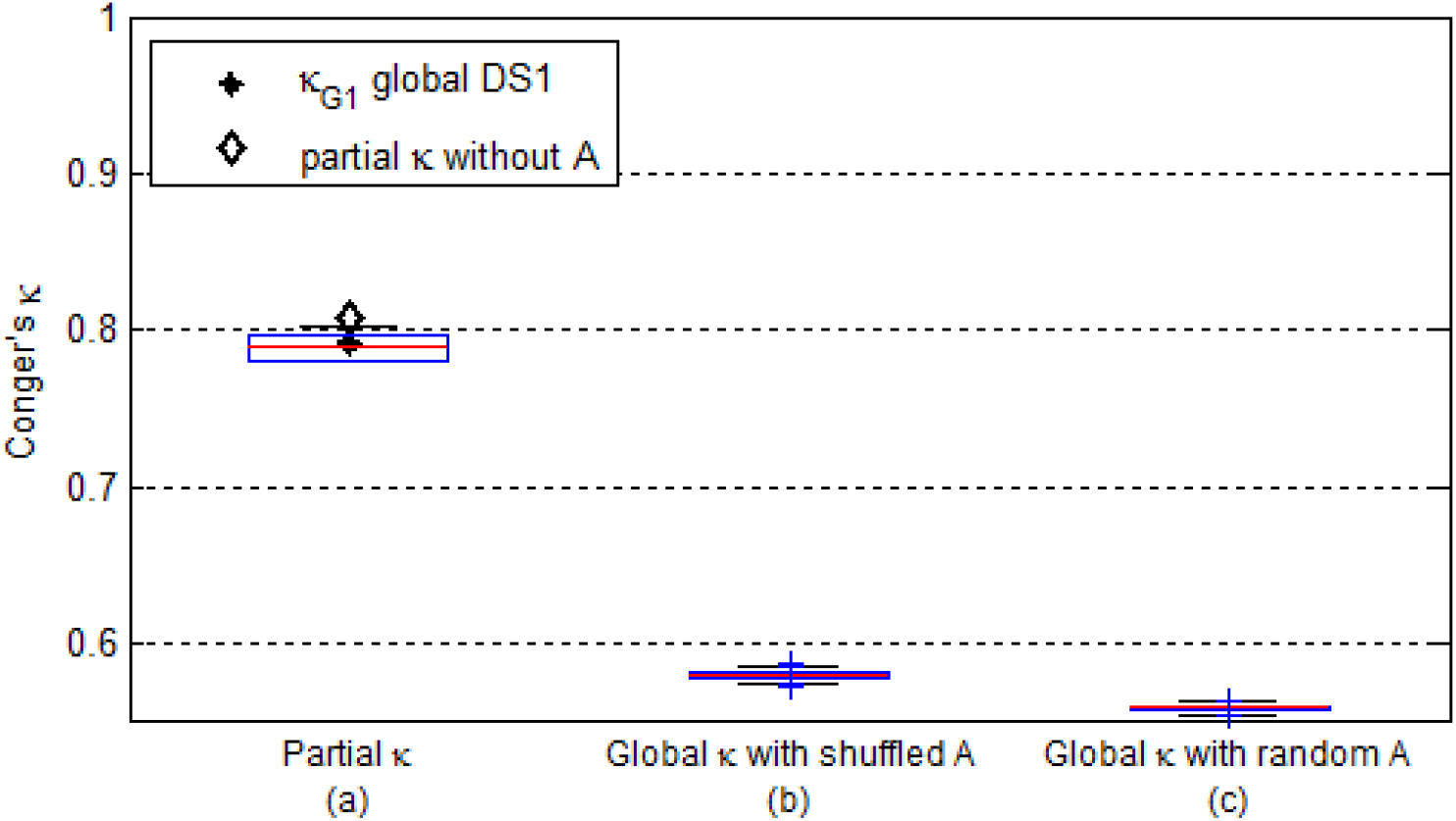
Influence of autoscoring. (a) Distribution of partial kappa coefficients when removing one scorer at a time by permutation, compared to the global kappa κ_G1_ (bold circle) that includes all scorings including the automatic one. The partial kappa coefficients ranged from 0.78 to 0.81 (μ=0.79). The specific case of “visual”partial kappa, computed without autoscoring, is highlighted in bold diamond. On the right, distributions of global κ coefficients when automatic scoring is substituted 1000 times (b) for a randomly shuffled scoring that preserves sleep stage prevalence ([0.57 − 0.59], μ=0.58) and (c) for a fully random scoring ([0.55 − 0.56], μ=0.56). Note that the good inter-expert agreement prevents the global kappa from reaching the chance agreement of 0 value when autoscoring is fully random.

In a fourth step, we compared scorings with various majority scorings V_Maj_. A high N_Maj_ was associated with a high agreement with V_Maj_, but implied a high number of rejected epochs for which too few experts agreed to reach a consensus (fig. 5). No expert obtained a 100% agreement with the majority scoring.

**Figure 5.**
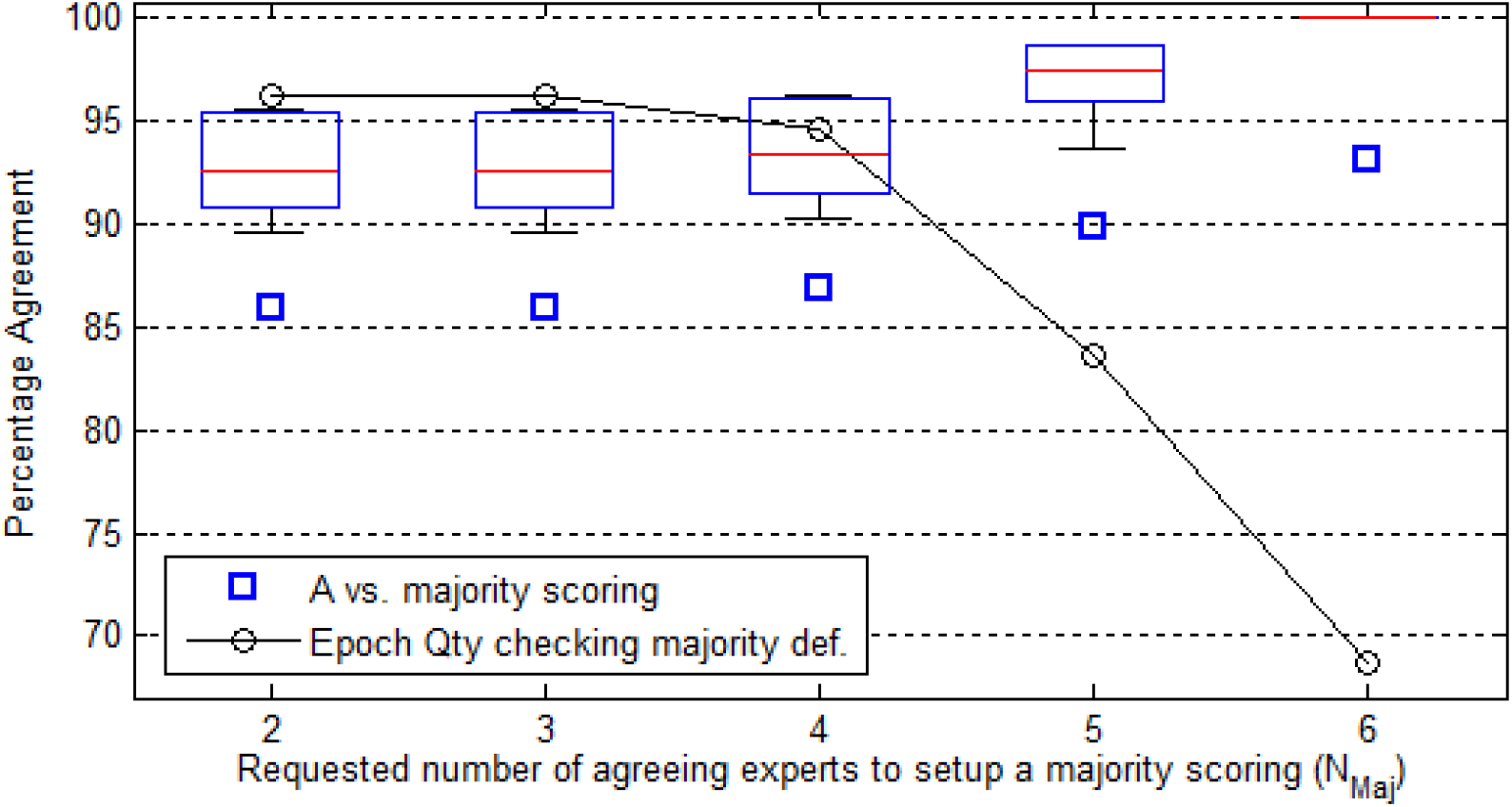
Majority scoring. Comparison between scorings and visual majority scoring, V_Maj_, used as the scoring reference. The evolution of the agreement between V_Maj_ and A (blue square) or V (boxplot) is drawn according to the minimal number of agreeing experts N_Maj_ requested to setup a majority. The number of valid epochs on which the comparisons are computed is also plot (black circle) according to N_Maj_. When more agreeing experts are requested to setup a majority, agreements increase but the number of valid epochs decreases. The last case, N_Maj_ = 6, equivalent to the visual full consensus V_all_, rejected a third of epochs. Importantly, the experts composing the majority scoring can be different at each epoch. For instance, when N_Maj_=3, the V vs. V_Maj_ agreement was [89.7% − 95.6%], μ=92.8% (86.0% for A vs. V_Maj_), 3.7% of epochs had ex-aequo majorities and 0.4% of epochs did not get 3 agreeing experts. So disagreements always exist between “real” visual scorings and “virtual” majority scoring, i.e. nobody scores like the majority scoring.

The subset of the most reliable epochs according to visual scoring was built by discarding the 1,349 epochs (31.3%) of disagreement between experts, leaving 2,959 epochs of full visual consensus. The contingency matrix is shown in Table 3. Automated scoring used as benchmark (1st column) shows that the highest disagreement among experts is observed for epochs autoscored N1, while higher agreement is observed among experts for Wake and N3.

**Table 3:**
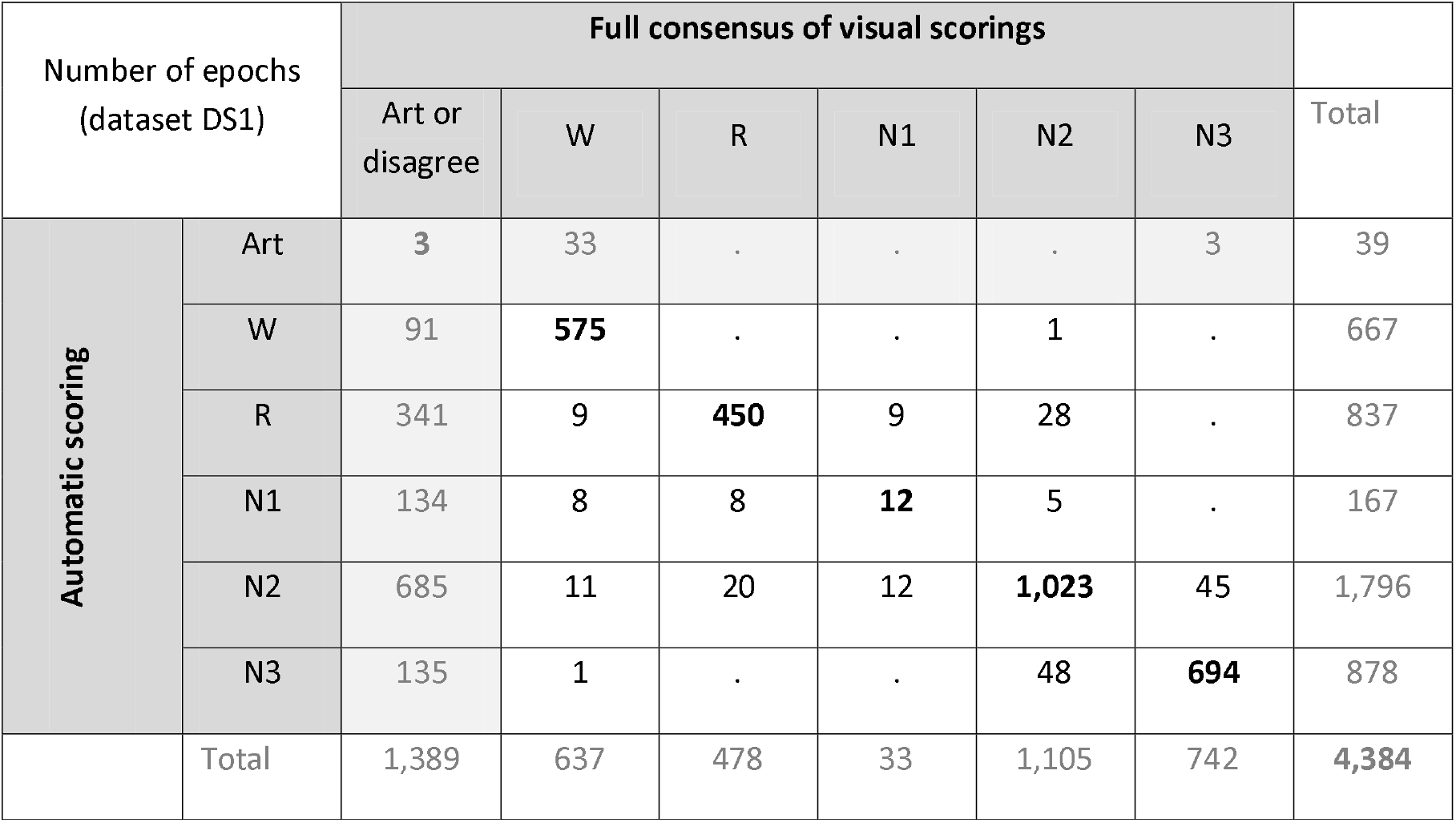
Contingency matrix for the dataset DS1. 1,389 epochs out of 4,384 have been discarded. 76 for artefact labeling by A or V, and 1,389 due to disagreement between visual scorers (1^st^ column).

Among these 1,349 non-consensual epochs between experts, Aseega agreed 1,255 times (93.0%) with at least one expert.

Proceeding from the contingency matrix, the Sensitivity and Positive Predictive Value of all sleep stages are reported in Table 4.

**Table 4:** Sensitivity (Se) and Positive Predictive Value (PPV) of automatic scoring vs. visual full-consensus (DS1). These results were computed after the removal of the 1,389 epochs of partial disagreement between experts.

| Dataset DS1 | W | R | N1 | N2 | N3 |
| --- | --- | --- | --- | --- | --- |
| Se | 95.2 | 94.1 | 36.4 | 92.6 | 93.9 |
| PPV | 99.8 | 90.7 | 36.4 | 92.1 | 93.4 |

### Dataset 2

Two recordings were rejected by Aseega because of low signal quality. Out of the remaining 102,141 recorded epochs, 764 (0.80%) were classified as artefact by automatic scoring and 445 (0.46%) were classified as artefact by the experts. 115 artefact epochs were common to automatic and visual scoring. The total number of discarded epochs was 1094 (1.14%), leaving 101,047 epochs for subsequent analysis.

Regarding the A vs. V comparison, the global kappa was κ_G2_ = 0.70 and the percentage agreement 79.0%. The 6 pairwise kappa coefficients between automatic scoring and each expert ranged [0.67 − 0.73], μ=0.70 (fig. 3a, right boxplot). The corresponding percentage agreements ranged [76.4% − 80.8%], μ=78.7 (fig. 3b, right boxplot).

Intra-expert variability across the two datasets using autoscoring as reference (step2) was assessed by pairwise agreements between DS1 and DS2. It systematically decreased from DS1 to DS2 for all experts (μ=-3.7%, fig. 6): kappa coefficients ranged [0.68 − 0.73], μ=0.70 and percentage agreements ranged [77.2 − 81.2], μ=79.0. Computation on reduced data showed that there was nearly no impact of having 20-fold less data in DS1 than in DS2.

**Figure 6.**
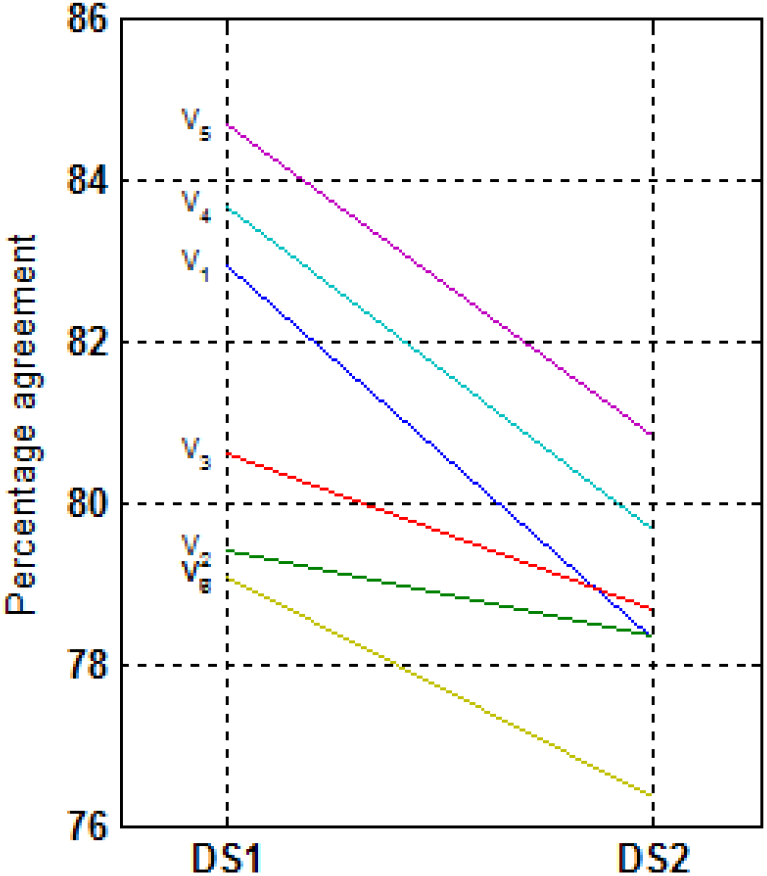
Intra-expert variability using autoscoring as reference. Impact of the scoring condition (temporal proximity of the dataset with the training sessions). For each visual scorer, V_i_, evolution of the percentage agreement with automatic scoring according to the dataset. The agreement decrease between datasets is not only a mean effect (μ=-3.7%), as illustrated in figure 3b, but also a systematic effect for every single expert, ranging from −1.3% to −5.6%.

Regarding intra-expert variability according to the sleep condition using autoscoring as reference, figure 7 shows that agreements on long duration nights (EXT and REC) were globally lower than the ones obtained on shorter duration nights (BAS and BEF).

**Figure 7.**
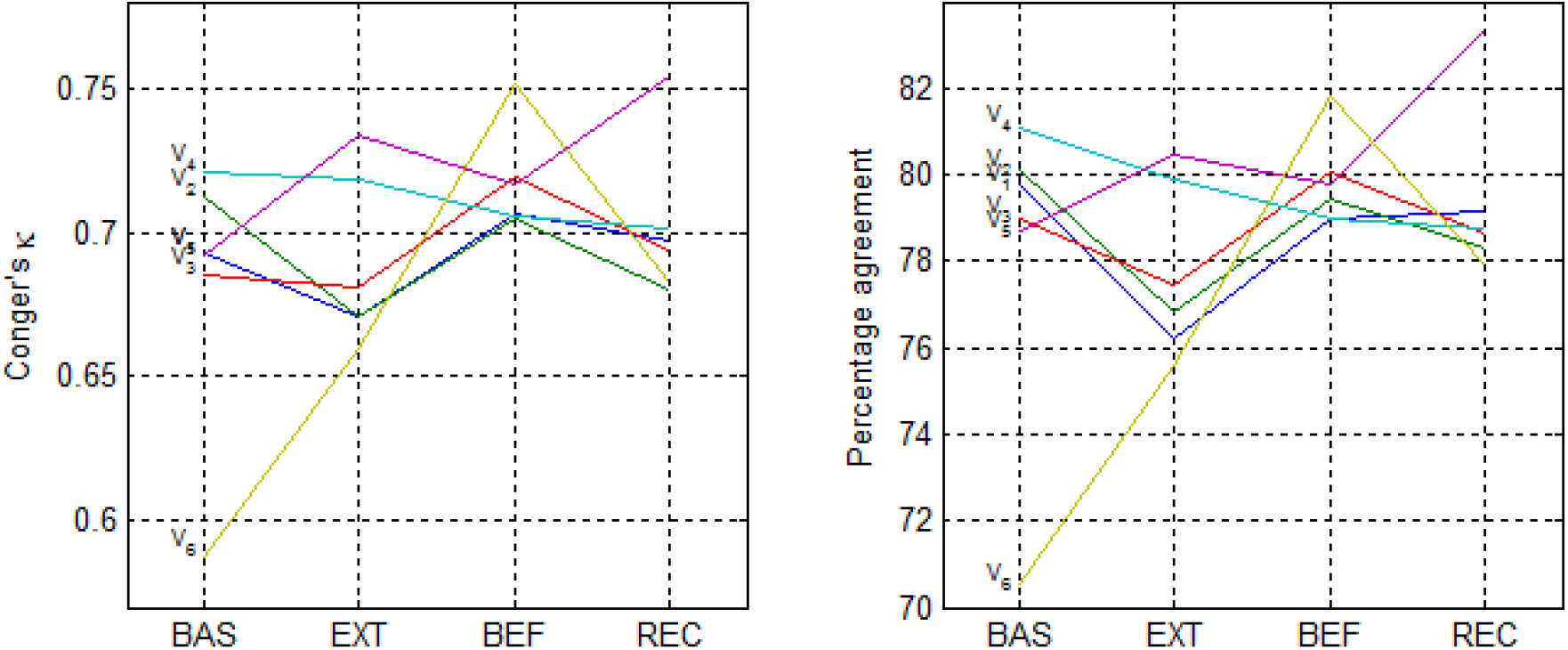
Intra-expert variability using autoscoring as reference. Impact of the sleep condition. For each visual scorer V_i_, agreements (Conger’s kappa on the left and percentage agreement on the right) are reported according to the sleep condition: baseline (BAS), extended night (EXT), before sleep deprivation (BEF) or recovery night (REC).

The contingency matrix of dataset DS2 (Table 5), provides the Sensitivity and Positive Predictive Value of all sleep stages (table 6).

**Table 5.** Contingency matrix for the dataset DS2. 764 epochs out of 102,141 have been discarded for artefact labeling by A and 445 by V.

| Number of epochs<br>(dataset DS2) |  | Visual scorings |  |  |  |  |  |  |
| --- | --- | --- | --- | --- | --- | --- | --- | --- |
|  |  | Art | W | R | N1 | N2 | N3 | Total |
| Automatic scoring | Art | 115 | 302 | 65 | 117 | 127 | 38 | 764 |
|  | W | 38 | 8,460 | 140 | 714 | 154 | 7 | 9,513 |
|  | R | 148 | 776 | 17,099 | 4,341 | 2,430 | 0 | 24,794 |
|  | N1 | 15 | 481 | 329 | 1,023 | 458 | 11 | 2,317 |
|  | N2 | 95 | 1,013 | 1,488 | 2,790 | 38,377 | 2,817 | 46,580 |
|  | N3 | 34 | 75 | 1 | 36 | 3,144 | 14,883 | 18,173 |
|  | Total | 445 | 11,107 | 19,122 | 9,021 | 44,690 | 17,756 | 102,141 |

**Table 6:** Sensitivity (Se) and Positive Predictive Value (PPV) of Automatic scoring vs. visual (DS2).

| Dataset DS2 | W | R | N1 | N2 | N3 |
| --- | --- | --- | --- | --- | --- |
| Se | 78.3 | 89.7 | 11.5 | 86.1 | 84.0 |
| PPV | 89.3 | 69.4 | 44.4 | 82.6 | 82.0 |

## Discussion

In this article, we assessed the reliability of sleep scoring, as conducted by 6 sleep experts and an automatic algorithm. The main results are threefold. First, a very good agreement is achieved between visual and automatic scoring, although automatic scoring is shown to marginally lower the overall scoring agreement. Second, agreement between experts decreases as the number of experts increases, suggesting that the issue of inter-expert variability is not restricted to a limited number of epochs that are difficult to score. Third, when referred to a reproducible method, such as autoscoring, visual scoring demonstrated intra-expert variability across datasets and type of recordings, thus introducing uncertainty in the measurement of the sleep variables of interest.

### Visual-automatic agreement is similar to visual-visual agreement

Overall, scoring agreement between scorers in DS1 was excellent^34^ (Conger’s kappa 0.79), while pairwise kappa between visual and automatic scoring ranged between 0.72 and 0.79, corresponding to a 82% agreement. The lowest agreement corresponded to the 6th expert, a new laboratory member under training to AAASM scoring rules. This may explain why this scorer obtained the lowest agreement with Aseega (fig. 6). The agreement increased to 93.1% when comparing automatic scoring with consensus scoring, due to the rejection of visually ambiguous epochs.

The best kappa coefficient was observed when automatic scoring was left out. Likewise, better pairwise kappa and percent agreement were observed between visual scorers (respectively, 0.81 and 86%). High agreement between experts from the same center guarantees high homogeneity in local scoring but does not rule out inter-site variability (not assessed here). Inter-site variability should be kept to a minimum.^4, 35^ Being marginally out of the pool of experts from a given sleep center allows the automated scoring to be transferrable to other sites.

A significant drop in automatic-visual agreement is observed between DS1 and DS2 for the REM PPV whereas the corresponding Se remains quite stable (Tab. 4 & 6). Qualitative appreciation of scorings showed that autoscoring tends to smooth REM episodes, unlike visual scoring where REM episodes are more fragmented. This fragmentation affects automatic-visual agreement when automatic is compared to one expert only. However, whereas commonly marked by all experts, this fragmentation is located differently by all experts. It therefore disappears from visual consensus scoring, and thus from the automatic-visual comparison. Here, being marginally different from a given expert allows the automated scoring to be transferrable to other expert. From another perspective, these results confirm that comparing any scoring, being automatic or visual, to a single expert is hazardous.

### Inter-expert variability - Visual disagreement is not just noise

As stated by Silber and colleagues when introducing the new AASM scoring rules: “no visual-based scoring system will ever be perfect, as all methods are limited by the physiology of the human eye and visual cortex, individual differences in scoring experience, and the ability to detect events viewed using a 30-second epoch”.^36^ In consequence, there is a continuous efforts to improve the guidelines (GRADE program in 2009^18^) and to homogenize their enforcement.^15^ Based on this program, a study including over 2,500 technicians from accredited laboratories showed 82.6% agreement compared to an epoch-by-epoch reference built as the majority score for each epoch.^15^ Even if this inter-scoring reliability may be overestimated, either for protocol reasons^1^ or by using majority scoring as the reference which can be misleading (see below), these continuous improvements in guidelines and training are needed and welcome.

In our study, when limiting the number of experts to 2, their percentage agreement is about 85%, whatever the pairing. However, this apparent homogeneity is misleading by suggesting that only 15% of epochs raise doubts. Increasing the number of experts significantly decreases the overall agreement and shows that the contentious epochs -and scoring consensus-depend strongly on which pair of experts is considered. On DS1, and for this team of sleep experts, we estimated that the asymptotic consensus agreement is closer to 65%, meaning that one third of the epochs raised doubts, in line with figures reported by Younes and colleagues.^22^

Our results show that the inter-expert disagreement cannot be considered as a low-level constant noise. It is not only a matter of specific epochs that are difficult to score, like transient or artefacted epochs.^22^ If so, adding more experts to build the scoring consensus would not affect the number of consensus epochs that way. We assume that the variability in inter-expert-agreement comes from both epoch-specific content that makes scoring rules difficult to apply, and expert-specific sensitivity to signal content (e.g. rapid or slow frequencies, sleep microstructures, etc.). In addition, we surmise that the inter-expert variability would further increase if we had considered clinical recordings with more altered sleep (e.g., sleep breathing disorders, periodic limb movement, and degenerative diseases) or experts from various sleep centers.^10, 12, 37^

The scoring consensus rejected a third of epochs. According to majority scoring and autoscoring, N1, N2 and R were the sleep stages most subject to disagreement between experts, whereas other studies found highest agreement for W, N2 and R.^15^ As stated above, the research-oriented visual scoring yielded REM episodes fragmented by multiple intrusions of N2 or N1 sleep stages, for all experts but located differently, thus increasing the disagreement for these stages.

Consensus scoring is a costly mitigation for inter-expert variability. First, because it requests at least 3 scorers to analyze the data; second, because it entails large amounts of rejected epochs hence of missing data. Majority scoring is not satisfactory either, for the same reasons. Besides, majority scoring suggests that a unified majority of agreeing experts exists, next to a minority of disagreeing experts. In reality, the composition of this majority changes at each epoch: no expert reaches 100% of agreement with the majority scoring (fig.5).

Among alternative approaches proposed in the literature to cope with the inter-expert variability, the “discussed consensus” implies that experts discuss on contentious epochs until they reach a consensus.^13, 19^ If this time-costly approach does reinstate the non-consensual epochs, inter-expert variability is abolished at the methodological cost of losing the independence of experts. Another alternative is the time- and cost-effective computer-assisted scoring, that has been explored since decades.^22, 38–40^ However it only copes partially with expert variability since it still involves human expertise.

### Intra-expert variability - Visual scoring is less reliable across datasets

Intra-expert variability, rooted in learning process, experience, and fatigue, is usually demonstrated by comparing different scorings performed on the same data by a given expert (score-rescore approach). However, even if this variability cannot be measured in the typical scoring routine (single-score), its effects still apply on each scored recording.

### Scoring conditions

Our results highlighted this intra-expert variability on different data by using autoscoring as the reference. The systematic decrease in visual-automatic agreement for each expert between DS1 and DS2 (fig. 6) originates from variability in visual scoring as the automatic scoring used in this study is fully deterministic and reproducible.^23^ The main assumption is a “drift” over time in visual scoring.^1^ As noticed by Danker-Hopfe and colleagues,^19^ training sessions represent an essential tool to achieve a homogeneous interpretation of the scoring rules. In our study, prior training sessions occurred right before DS1 scoring, which was then followed by DS2 scoring. This might explain why we observe an excellent inter-scorer agreement for the DS1. In contrast, DS2 scoring seemed to reveal the individual drift, away from common scoring rules. This drifting over time generates both inter and intra expert variability and is a critical concern in clinical and epidemiological research, and specifically in our current focus on scoring massive datasets.

### Sleep conditions

Long nights (EXT and REC) showed globally lower agreements compared to shorter ones (fig. 7), suggesting that the intra-expert variability is possibly determined by fluctuations in attention. In addition, the intrinsic composition of each night appears to matter. For instance, the REC night which expectedly showed more consolidated sleep gives rise to a better agreement between automatic and visual scoring. By contrast, the EXT night, which is associated with more fragmented sleep (i.e. more transitions) at the end of the night due to the decreased sleep debt, yields to a lower agreement with a wider dispersion of these agreements.

The 6^th^ (junior) expert showed a particularly low agreement with Aseega for the BAS night of DS2 (fig. 7) compared to the others. Actually, the BAS night is usually used as a reference for the subsequent recordings to be scored, where the expert “learns” to recognize the specific sleep patterns of a given participant. We hypothesized that this participant-specific learning process is slower for the junior expert than for senior experts, for whom the first few sleep cycles are sufficient.

### Automatic scoring: pros and cons

The investigated variability of sleep characteristics, *V_characteristic_*, is only reachable via analysis methods that introduce additional noise *N_method_*, such that the variability of the measurement *V_observed_* writes:

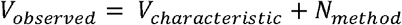

What is at stake is to minimize as much as possible the contribution of the noise of the method. By neutralizing external sources of variance such as inter and intra-expert variability, autoscoring avoids the possible masking effect of visual scoring, and gives closer access to the intrinsic meaningful variance of the data – even when it is of low amplitude. One could think of the interesting challenge of re-opening the cold cases of non-conclusive studies by re-analyzing data with autoscoring.

However, reproducibility safeguards from score-rescore variability, not from errors. Indeed, a systematic decision making such as autoscoring’s can be systematically wrong. There are two nonexclusive ways to address this major issue.

First, the use of autoscoring should be responsible and imply a systematic strategy of post analysis quality check: first, in order to maintain a close link to the data, second in order to identify and handle bad outputs. Bad outputs could be either discarded or corrected. Correcting bad output is an option proposed by some autoscoring services, or it can also be the expert’s task. Data are given a second chance, but variability, inherent to any visual operation, kicks back in: to what extent, for what effect, this should be assessed for acceptability on a case basis. Discarding bad outputs is more stringent, increases the amount of missing data, but the remaining data are 100% consistent.

Second, autoscoring methods should be precisely evaluated to identify potential limits. The result of the Aseega algorithm in this study confirmed the general trend towards a dramatic improvement in methods that long suffered from questionable performances and poor reputation. A number of automatic methods are currently available, yet, precise assessment and comparison on common datasets using common metrics^1, 2, 13, 41^ is an open question: the issue of inter-scorer variability is transferred among autoscoring methods. These methods differ in nature or by their objective: some are based on multichannel PSG data analysis,^42, 44^ others use a single EEG channel only^23, 45, 46^ or EOG only,^47^ some are limited to Wake-Sleep scoring.^48^ This should not necessarily affect expectations in terms of reliability. Published methods also differ by the way they are assessed,^39, 42–46, 49, 50^ study protocol (population studied and number of experts) and comparison methodology (reference setup and statistics). Providing public sleep databases is an ongoing and useful process for several years in the USA (www.sleepdata.org^35^), in Europe (www.physionet.org^51^) and more recently in Canada (www.ceams-carsm.ca/en/MASS^52^). It will be challenging to provide data fulfilling the requirements of all methods and … satisfactory scoring reference for these data, as demonstrated in this paper. Our contribution to this open database policy will be to open-source the data used in this study, including both visual and autoscoring.

## Acknowledgments

This study was funded by the Walloon Excellence in Life sciences and Biotechnology (WELBIO), Belgium. The authors are grateful to Erik Lambot, Audrey Golabek and Muriel Lennertz for their assistance in data collection.

## Abbreviations

EEG: – Electroencephalogram
EOG: – Electro-oculogram
EMG: – Electromyogram
PSG: – Polysomnography
BAS: – Baseline night
EXT: – Extended sleep opportunity
BEF: – Night before sleep deprivation
REC: – Recovery night after sleep deprivation
V: – visual scorer
A: – automated analysis Aseega
DS1: – Dataset 1
DS2: – Dataset 2

